# Recombinant reporter phage rTUN1::*nLuc* enables rapid detection and real-time antibiotic susceptibility testing of *Klebsiella pneumoniae* K64 strains

**DOI:** 10.1101/2022.08.19.504497

**Authors:** Peter Braun, Rene Raab, Joachim J Bugert, Simone Eckstein

**Affiliations:** Bundeswehr Institute of Microbiology, Munich, Germany

## Abstract

The emergence of multi drug resistant (MDR) *Klebsiella pneumoniae* (*Kp*) strains constitutes an enormous threat to global health as MDR associated treatment failure causes high mortality rates in nosocomial infections. Rapid pathogen detection and antibiotic resistance screening is therefore crucial for successful therapy and thus, patient survival. Reporter phage-based diagnostics offer a way to speed up pathogen identification and resistance testing, as integration of reporter genes into highly specific phages allow real-time detection of phage replication and thus, living host cells. *Kp* specific phages use the host’s capsule, a major virulence factor of *Kp*, as receptor for adsorption. To date, 80 different *Kp* capsule types (K-serotypes) have been described with predominant capsule types varying between different countries and continents. Therefore, reporter phages need to be customized according to the locally prevailing variants. Recently, we described the autographivirus vB_KpP_TUN1 (TUN1), which specifically infects *Kp* K64 strains, the most predominant capsule type at the military hospital in Tunis (MHT) that is also associated with high mortality rates. In this work, we developed the highly specific recombinant reporter phage rTUN1::*nLuc*, which produces Nanoluciferase (nLuc) upon host infection and thus, enables rapid detection of *Kp* K64 cells in clinical matrices such as blood and urine. At the same time, rTUN1::*nLuc* allows for rapid antibiotic susceptibility testing and therefore identification of suitable antibiotic treatment in less than 3 hours.

## Introduction

The ESKAPEE bacteria (*Enterococcus faecium*, *Staphylococcus aureus*, *Klebsiella pneumoniae*, *Acinetobacter baumanni*, *Pseudomonas aeruginosa*, *Escherichia coli* and *Enterobacter* species) are responsible for the majority of nosocomial infections and typically possess multi-drug resistance (MDR). Among these, the opportunistic pathogen *K. pneumoniae* (*Kp*) poses a particular threat for public health. Ubiquitously occurring as a commensal, *Kp* can be found among gastro-intestinal microbiota, in the respiratory tract and on the skin. As a facultative pathogen, *Kp* can cause a variety of severe nosocomial diseases such as pneumonia, sepsis, wound- and urinary tract infections (UTIs). In recent years, the number of community-acquired cases of *Kp*-caused pneumonia and meningitis has dramatically increased (Herridge et al., 2020). Together with the emergence of acquired MDR against broad-spectrum antibiotics (ABs) and the natural resistances of the bacterium, treatment of *Kp* infections has become a global healthcare challenge (Babini & Livermore, 2000; Knothe et al., 1983; Prestinaci et al., 2015).

Infections with carbapenem-resistant *Kp* are associated with serious symptoms and high mortality rates (Patel et al., 2008; Schwaber et al., 2008). Therefore, rapid diagnosis of the infection as well as detection of resistance markers are crucial for patient survival. Classical approaches to identify carbapenem-resistant *Kp* rely on bacterial cell culture with subsequent biochemical profiling and therefore provide results only after a few days. While nucleic acid based tools such as diagnostic real-time PCR or loop-mediated isothermal amplification (LAMP) enable fast and specific detection of the pathogen, the established assays mainly target carbapenem-resistance genes and therefore cannot distinguish between living cells and DNA residues present in the sample (Hartman et al., 2009; Kurupati et al., 2004; Nakano et al., 2015).

Diagnostic bacteriophages (phages) represent a promising alternative approach. In addition to their therapeutic potential, phages are suited for the identification of bacteria due to the viruses’ high specificity. Phage sensitivity assays have been in practice for centuries for a variety of pathogens such as *Bacillus anthracis* (Turnbull et al., 2008) or *Yersinia pestis* (Garcia et al., 2008). Such techniques include antibody-based detection of phages using ELISA (Stewart et al., 2013) or detection of amplified phage DNA (Anany et al., 2018). Another option is to exploit the bacteriolytic activity of phages and detect released cellular components such as ATP (Blasco et al., 1998). The most promising methodology, however, is the use of reporter phages, which either have a reporter molecule directly attached to the virion surface or produced during phage replication (Goodridge et al., 1999; Hennes & Simon, 1995; Meile et al., 2020).

While most phages of gram negative bacteria utilize surface structures such as lipopolysaccharides (LPS), teichoic acids or proteins as host receptors for adsorption (Nobrega et al., 2018), the specificity of *Kp* phages is mostly determined by the different *Kp* capsule polysaccharides (CPS). CPS represent a major virulence factor of *Kp* as the thick layer protects the bacterial cell from phagocytosis and prevents complement binding during host response. In addition, the CPS forms a physical barrier towards ABs as it hampers diffusion into the bacterial cell (Domenico et al., 1994; Merino et al., 1992; Shon et al., 2013). To date, about 80 different *Kp* capsule serotypes (K-types) have been described (Pan et al., 2015). Since *Kp* specific phages are usually capable of infecting only one or a few capsule types, the selection of a suitable reporter phage is crucial for reliable diagnostics. As predominant capsule types vary from region to region, reporter phages need to be customized according to the locally prevailing variant. In Tunisia, e.g., the majority of nosocomial infections at the Military Hospital of Instructions in Tunis (MHT) are caused by a *Kp* strain featuring capsule type 64 (K64). These *Kp* K64 infections are associated with high mortality rates, especially among intensive care patients (Eckstein et al., 2021).

In this study, we genetically engineered the recently identified *Kp* K64-specific phage vB_KpP_TUN1 (TUN1), and constructed a recombinant TUN1 reporter phage (rTUN1::*nLuc*) for the rapid detection of *Kp* K64 strains in bacterial cell culture or directly from clinically relevant matrices as well as for conducting real-time antibiotic susceptibility testing.

## Material and Methods

### Bacterial growth

All *K. pneumoniae* and *E. coli* strains were cultivated in LB-broth [10% NaCl (w/v), 10% peptone (w/v), 5% yeast extract (w/v)] at 37° C under constant shaking.

### In vitro DNA-Assembly of synthetic phage genomes

For isolation of genomic DNA (gDNA) of phages Master-Pure™ Complete DNA and RNA Purification Kit (Lucigen, Wisconsin, USA) was used according to the supplier’s protocol. Prior to assemble the synthetic TUN1 WT genome, TUN1 WT gDNA was used as PCR template to amplify five fragments. This was achieved by using the primer pairs TUN1 F1 + R1 (= Fragment 1), TUN1 F2 + R2 (= Fragment 2), TUN1 F3 + R3 (= Fragment 3), TUN1 F4 + R4 (= Fragment 4) and TUN1 F5 + R5 (= Fragment 5). Primers were designed to generate 15-25 bp overlaps between adjacent fragments to ensure efficient assembly. The circular approach was done in the same way, with the difference of replacing Fragment 1 and Fragment 5 by Fragment 5_1, amplified using the primers TUN1 F5 + R1 (Supplementary Figure S1A).

If not stated differently, the four TUN1 fragments of the circular approach described above were used for all further TUN1 DNA assemblies.

In order to generate the deletion strain TUN1 Δhpgc1, Fragment 5_1 was replaced by Fragment 5_1 gp1 (using the primer pair TUN1 F5 + gp1 R) and Fragment 2 by Fragment 2 gp5 (primers TUN1 gp5 F and 2R). TUN1 Δhpgc2 was achieved by using Fragment 5_1 gp6 (primers TUN1 F5 and gp6 R) and Fragment 2 gp9 (primer pair TUN1 gp9 F + R2). For TUN1 Δhpgc3 construction, the primer TUN1 gp9 R and gp12 F were used instead of primer TUN1 2R and 3F, respectively, resulting in the new amplicons Fragment 2 gp9_2 and Fragment 3 gp12 (Supplementary Figure S1D). In each case, TUN1 WT gDNA was used as a PCR template.

To generate the triple mutant TUN1 Δhpgc123, Fragment 3 and Fragment 4 were amplified from TUN1 WT gDNA using the primer pairs mentioned above. Amplification of modified Fragment 1 was achieved by using the primers TUN1 F5 and gp6 R and gDNA of TUN1 Δhpgc1 as a PCR template. Furthermore, Fragment 2 was amplified from TUN1 Δhpgc2 gDNA by using the primer pair TUN1 gp9 F + R2 (Supplementary Figure S1E).

To construct rTUN1::*nLuc* either TUN1 WT or TUN1 Δ*gp7-8* gDNA was used as a PCR template. In both cases, four fragments (F 5_1, 2, 3 and 4) were amplified using the primers TUN1 fwd5 + TUN1 rev1, TUN1 fwd2+ rev2, TUN1 fwd3 + TUN1 MajCap rev, TUN1 MinCap fwd + TUN1 rev4. *nLuc* was amplified from pNL2.1 (Promega, Walldorf, Germany) using the primer pair nLuc IGR Cap fwd/rev.

For all approaches DNA-Assembly was conducted using the NEBuilder DNA-Assembly Master Mix (New England Biolabs, Ipswich, USA) according to the manufacturer’s protocol. Oligonucleotide sequences are listed in Supplementary Table S1. Primer design and *in-silico* cloning was carried out using Geneious Prime (Version 2021.1.1, Biomatters New Zealand)

### Phage rebooting

Phage assembly and rebooting was achieved in a non-replicative host by transforming chemically competent *E. coli* 10-beta cells (New England Biolabs) with 5 μl of the assembly mix according to the protocol provided by the manufacturer. After transformation 950 μl of LB medium were added and the cells grown for 2 hours at 37° C with horizontal shaking at 120 rpm. Subsequently, the cells were harvested for 5 min at 3,000 × g and the supernatant, containing rebooted phages, stored at 4° C until further use.

### Plaque-Assay

To check for functionality of (recombinant) TUN1 a fresh culture of *Kp* was grown to OD_600_= 0.4-0.6 and then 350 μl of the culture were mixed with 2.5 ml hand-warm soft-agar [LB + 0.6 % Agar (w/v)] together with 100 μl of phage stock/ transformation supernatant. Then, the mixture was evenly distributed on a LB-agar plate and, after solidification, the plates were incubated at 37° C for 16 h. The next day, plaque forming units (PFUs) were calculated and plaque morphology analyzed.

### Bacterial growth

To determine the bacteriolytic activity of TUN1 phages on *Kp* strains, bacterial growth was measured comparing samples with and without phages. For this purpose, 100 μl of the respective *Kp* culture (10^7^ CFUs/ml) were mixed with either 100 μl of TUN1 phage stock (1×10^3^ PFUs/ml) or 100 μl of LB medium as a control in a 96-well plate with clear bottom. Plates were incubated at 37° C with shaking and bacterial growth (OD_600nm_) was measured every 30 min.

### rTUN1::*nLuc* functionality

To measure luciferase activity on plaque containing LB-agar plates, 5 μl of the diluted (1:50, in PBS) NanoLuc^®^ luciferase substrate 2-furyl methyl-deoxy-coelenterazine (Furimazine; Promega) were dropped onto the respective plaques. After incubation at RT for 3 min, the plates were analyzed for luminescence signals using a ChemiDoc Imaging System and Image Lab 6.0.1 software (Bio-Rad Laboratories Inc., Hercules, USA). Exposure time was set to 5 sec. The obtained luminescence signals were then merged with an image of the plate for identification of plaques featuring luciferase activity.

To analyze luminescence in liquid culture medium over time, 100 μl of a fresh *Kp* culture (1×10^7^ – 1 CFUs/ml) mixed with 100 μl of rTUN1::*nLuc* stock (1×10^3^ PFUs/ml) were transferred into a cavity of a 96-well plate and supplemented with 2 μl of substrate. The plate was then incubated at 37° C under continuous shaking (420 rpm) using a Varioskan Lux plate reader (Thermo Fisher Scientific, Waltham, USA). Bacterial growth (OD_600nm_) and luminescence (exposure time: 100 ms) were measured every 30 min for up to 16 h.

### Clinical matrices

For rTUN1::*nLuc* based *Kp* detection in clinically relevant matrices, the reporter phage was added to *Kp*-spiked urine and blood samples. Fresh urine was sterile filtered (0.22 μm luer lock syringe filter, Merck Millipore, Cork, Ireland) spiked with the TUN1 host *Kp* 7984 (final conc. 1×10^7^ CFUs/ml). Subsequently, bacterial growth and luminescence were measured in a plate reader every 30 min for 7 h at 37° C as described above.

To test the clinical matrix blood, 5 ml defibrinated sheep blood, spiked with the *Kp* 7984 (final conc. 1×10^7^ CFUs/ml) was used. Prior to measurements, the serum was collected as described elsewhere (Matsui et al., 2019). Briefly, the 5 ml spiked blood were injected into a 50 S-Monovette^®^ (Sarstedt, Nuernbrecht, Germany) and centrifuged at 2,000 × g for 15 min. The serum containing the bacterial cells was then transferred into a fresh tube and centrifuged again. After discarding the supernatant, the bacteria were resuspended to the original volume using LB medium. Subsequently, bacterial growth and luminescence with and without the presence of rTUN1::*nLuc* was measured as described above.

### Data analysis

Sequence data was analyzed using Geneious Prime (Version 2021.1.1). All Varioskan data was analyzed using the SkanIt 6.1 RE for Microplate Readers RE Software (Version 6.1.0.51) and GraphPad Prism (Version 8.4.3).

## Results

### Successful rebooting of TUN1 from synthetic phage genomes requires circularized DNA

Generation of TUN1 (an Autographivirus featuring a 41 kb sized genome) reporter phages was aimed to be achieved by synthetic biology driven genetic engineering. As a proof of concept, the technical feasibility of *in vitro* assembly and phage rebooting had to be demonstrated prior to construction of reporter phages. For this, the TUN1 wildtype (WT) DNA was PCR amplified as five or four overlapping fragments and assembled *in vitro* generating linear or circularized synthetic phage genomes, respectively. Recombinant phage TUN1 was successfully rebooted by transforming the non-replicative host *E. coli* NEB Stable with circularized synthetic constructs while transformation of linear constructs did not result in any rebooted phages (Supplementary Figure S2). In a next step, we aimed at integrating the NanoLuc^®^ luciferase gene (*nLuc*) fused to a ribosomal binding site (RBS) into the synthetic circular construct. As insertion site for the RBS-*nLuc* construct, the intergenic region between the major (*gp38*) and the minor (*gp39*) capsid genes was selected (Figure 1), as it qualifies for efficient transcription of the reporter due to the strong *cps* promoter regulating this operon (Loessner et al., 1996a). After *in vitro* DNA-assembly and phage rebooting, however, no plaques were observed on the *Kp* K64 strain 7984 indicating that no functional recombinant reporter phage particles had been generated. Thus, while recombinant phage can be assembled from DNA fragments and rebooted, a recombinant phage with the additional reporter gene *nLuc* cannot.

**Figure 1.**
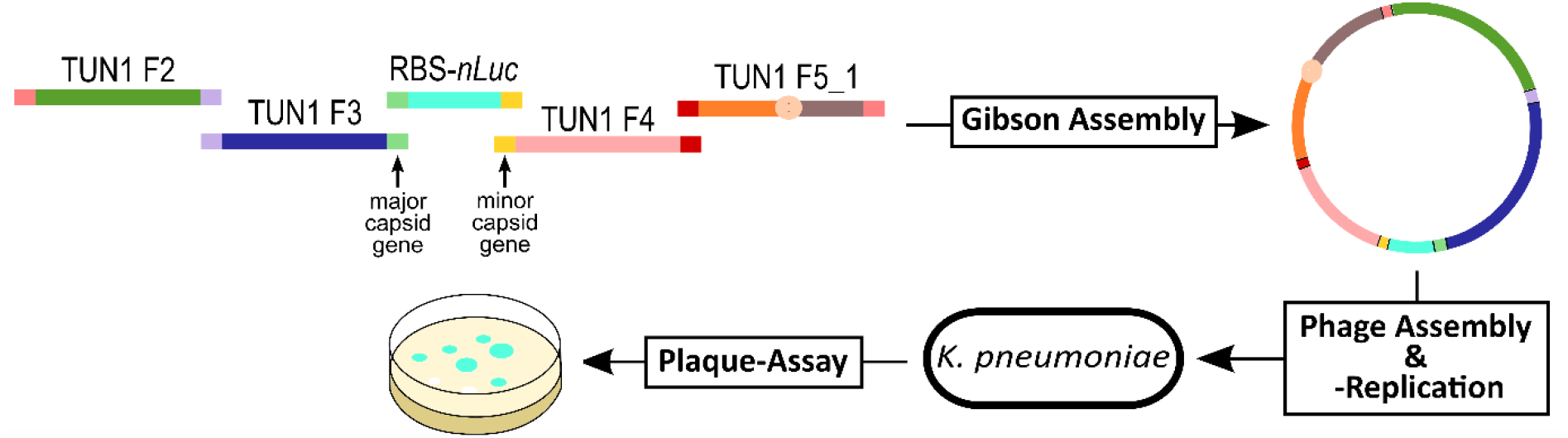
Schematic overview of TUN1 repoter phage construction. For construction of a TUN1 reporter phage all synthetic DNA fragments were amplified with overlapping ends between consecutive segments (depicted in matching colors). After *in vitro* DNA assembly the circular DNA was used to transform *E. coli* cells in which the recombinant phages were rebooted. Cells were lysed to release virions from the phage insusceptible host and the lysate used to infect *Kp*. Next, recombinant phage was *in vivo* amplified in its host *Kp* forming plaques on a lawn in the bacteria. Functional reporter phages were screened by measuring plaque luminescence after addition of luminogenic substrate.

### Genome reduction of TUN1 enables insertion of nLuc-reporter

One possible explanation for unsuccessful reporter integration was that the capacity of DNA encapsulation in the nascent virions is already reached with the size of the WT genome of TUN1. To address this challenge, we next aimed to reduce the TUN1 WT genome to provide space for insertion of the *nLuc* construct. As TUN1 comprises multiple ORFs encoding proteins of unknown function (Eckstein et al., 2021), these hypothetical genes (hpg) without any known function represent ideal candidates for targeted gene deletion. We decided to eliminate the operons *gp2-4*, *gp7-8* and *gp10-11* (Figure 2A) as these clusters were predicted to be regulated by single promoters, respectively. For this, we used the same method as described for TUN1 WT *in vitro* assembly above but excluded the selected hpg clusters (hpgc) during fragment amplification (Supplementary Figure S1), resulting in TUN1 Δhpgc1, Δhpgc2 and Δhpgc3 featuring genome reductions by 518 bp, 529 bp and 738 bp, respectively (Figure 2B). As all hpgc deletion variants resulted in functional phages, a triple deletion strain (TUN1 Δhpgc123) was generated by using the single-cluster deletion mutants as PCR-templates for subsequent synthetic phage genome rebooting. This resulted in a reduced genome size by 1,785 bp (Figure 2B). Analysis of the recombinant phages for their lytic activities against their host *Kp* 7984 in growth experiments, yielded no difference between TUN1 WT and the deletion variants (Figure 2C). While the plaque sizes of TUN1 Δhpgc1 and Δhpgc2 on *Kp* 7984 were similar to that of TUN1 WT, those of the deletion variants TUN1 Δhpgc3 and Δhpgc123 were slightly decreased. As the size of genome reduction in TUN1 Δhpgc2 (529 bp) perfectly matched the size of the *nLuc* construct (526 bp) this variant was used for reporter gene insertion in the next step.

**Figure 2.**
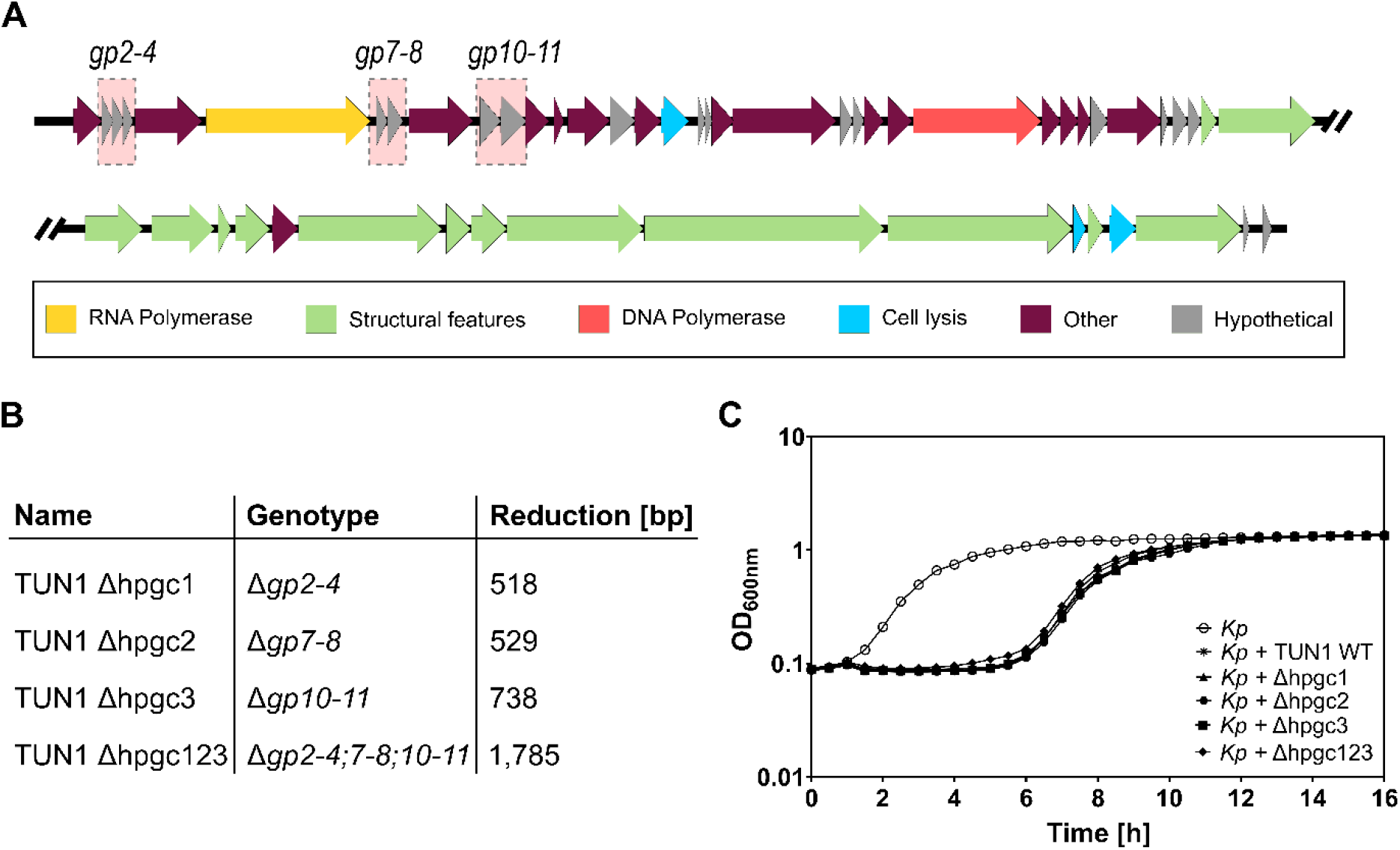
Characteristics of TUN1 deletion variants. **A)** Schematic illustration of TUN1 genome. Hypothetical gene clusters *gp2-4*, *gp7-8* and *gp10-11* are shaded in red. **B)** TUN1 deletion variants and their respective genome reduction. **C)** Growth curves of *Kp* 7984 without phage (○), in the presence of TUN1 WT (✱), Δhpgc1 (▲), Δhpgc2 (●), Δhpgc3 (■) or Δhpgc123 (◆).

Using the genome-reduced TUN1 Δhpgc2 as a template, we were able to insert the *nLuc* construct and thus to generate phage rTUN1::*nLuc*. Just like TUN1 WT and TUN1 Δhpgc2, rTUN1::*nLuc* also formed distinct clear plaques surrounded by translucent halos when tested in plaque assay on *Kp* 7984. To ascertain the functionality of the luciferase reporter, rTUN1::*nLuc* plaques were spot-covered with furimazin, the substrate of NanoLuc^®^ luciferase, and subsequently analyzed for luminescence. Indeed, all plaques emitted light upon addition of furimazin (Figure 3) and therefore, rTUN1::*nLuc* can be used for luminescence-based detection of *Kp* K64.

**Figure 3.**
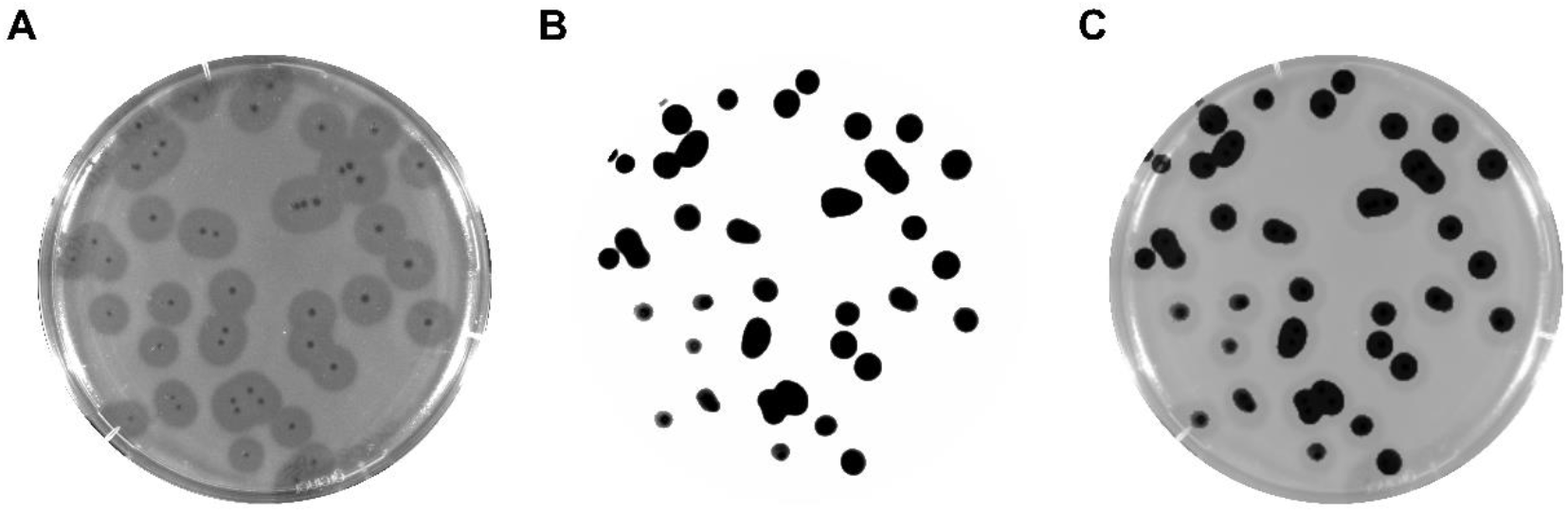
Nanoluciferase activity of rTUN1⸬ *nLuc* plaques. **A**) Photographic image of a typical plaque assay agar plate with recombinant reporter phage rTUN1::*nLuc* on *Kp* 7984 cells. **B**) Luminescent signal emitted from the agar plate (A) after addition of luminogenic substrate (exposure time: 5 sec). **C**) Merged image of A and B.

### rTUN1::*nLuc* enables highly sensitive *Kp* K64 detection

Plaque assays depend on bacterial growth on agar plates and are thus too time consuming for *Kp* detection in clinical diagnostics. Therefore, the next step was to test the applicability of the reporter phage rTUN1::*nLuc* for rapid luminescence-based *Kp* detection in liquid culture. For this, a liquid culture of *Kp* 7984 (1×10^6^ CFUs per well) was infected with 1×10^2^ PFUs (per well) of the reporter phage and bacterial growth and luminescence were measured over time. We observed an immediate strong increase of luminescence reaching a maximum of 10^8^ relative light units (RLUs) after only 2 h of incubation. This increase in luminescence correlated well with the start of impaired bacterial growth compared to the control containing only *Kp* without rTUN1*::nLuc* (Figure 4A). Most probably caused by autodegradation of the substrate, some luminescent background noise was also detected for *Kp* 7984 without phage. Therefore, the threshold for positive luminescence results was set to RLUs ≥ 10^3^. When different rTUN1::*nLuc* titers were tested, we observed high baseline luminescence (≥ 10^3^ RLUs) for phage titers greater than 1×10^3^ PFUs/well, even in phage-only control samples. This is most probably due to carryover of NanoLuc^®^ luciferase during preparation of rTUN1::*nLuc* phage stocks. An initial titer of 1×10^2^ PFUs/well, on the other hand qualified as ideal phage concentration as the baseline was comparable to the one produced by the no-phage control (Supplementary Figure S3).

**Figure 4.**
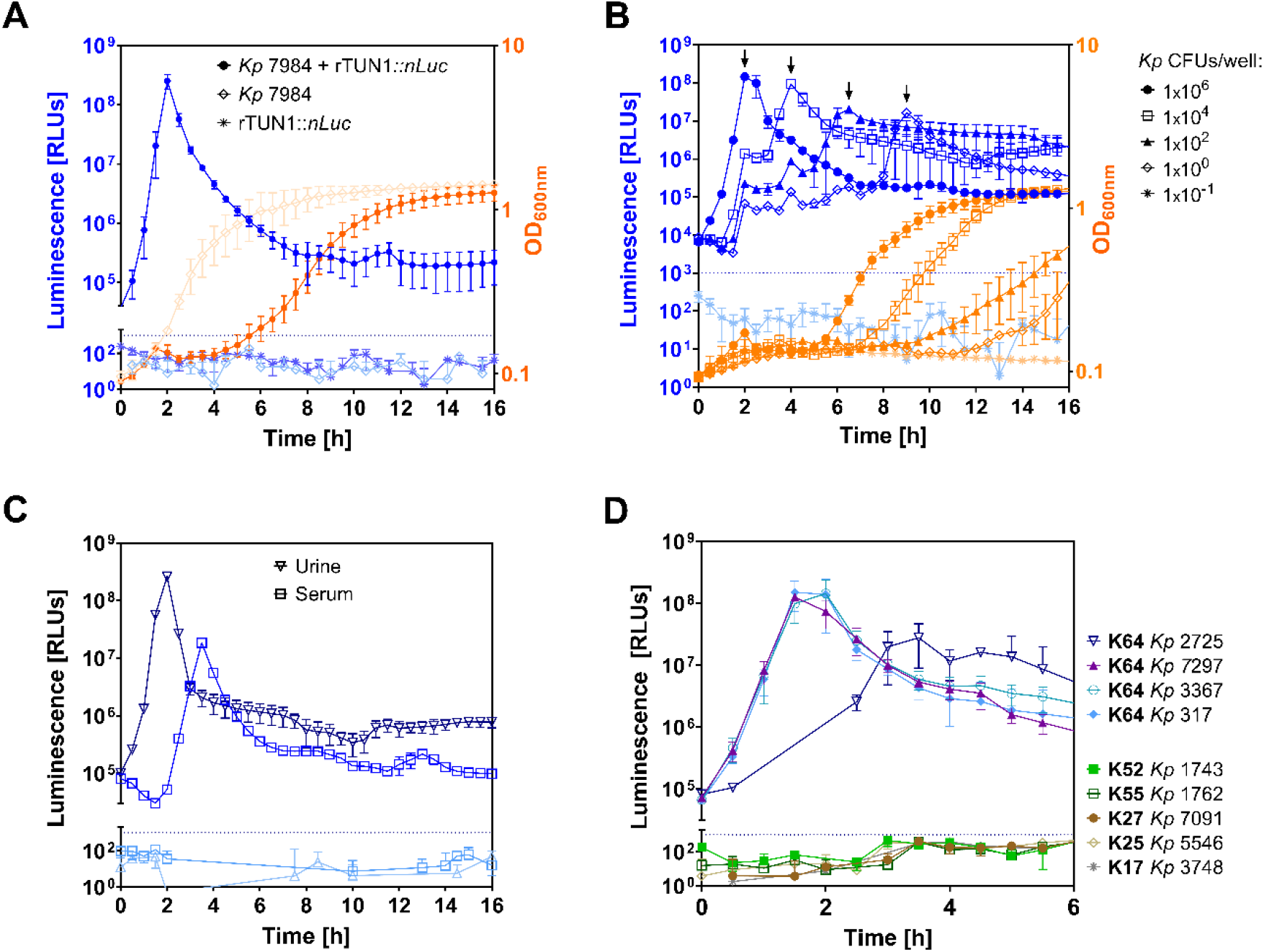
Luminescence based detection of *Kp* K64 using rTUN1⸬ *nLuc*. **A)** Luminescence (blue) and bacterial growth (orange) measured for *Kp* 7984 (1×106 CFUs/well with (●) and without rTUN1::*nLuc* (◇) as well as solely rTUN1::*nLuc* (✱) over time. **B)** Luminescence (blue) and bacterial growth (orange) of different *Kp* 7984 concentrations per well (●: 1×10^6^ CFUs; **□**: 1×10^4^ CFUs; *▲*: 1×10^2^ CFUs; ◇: 1×10° CFU; ✱: 1×10^−1^ CFU) in the presence of rTUN1::*nLuc*. Black arrows (⬇) indicate the luminescence peaks generated by rTUN1::*nLuc* on respective bacterial culture. **C)** Luminescence for *Kp* 7984 + rTUN1::*nLuc* (dark blue) and without phage (light blue) in urine (▽) and serum (**□**). **D)** Luminescence of rTUN1::*nLuc* with *Kp* of different K-types (K64, K52, K55, K27, K25 and K17). For every experiment, an initial titer of 10^2^ PFUs per well of the respective phage was used. Dotted blue line = threshold of 10^3^ RLUs.

To investigate the sensitivity and thus the detection limit of the rTUN1::*nLuc* based reporter phage assay, a dilution series of *Kp* 7984 (1×10^6^ – 0.1 CFUs per well) was tested using a starting concentration of 1×10^2^ PFUs/well of the reporter phage. Both the increase in detectable bacterial growth as well as the luminescence signal peak was delayed by 2-4 h with every dilution step. (Figure 4B). From this data a standard curve was generated which can be used to easily determine *Kp* concentrations based on the time of luminescence peak (Supplementary Figure S4). The results revealed that as few as one initial cell per well was sufficient to yield a luminescent signal with its peak at 10^7^ RLUs after approx. 9 hours. However, some of these wells with 10° *Kp* cells turned out negative (in luminescence and optical density) most probably because random distribution of single cells results in empty wells in some cases.

### *Kp* K64 cells can be detected directly in clinical matrices using rTUN1::*nLuc*

*Kp* can cause severe infections with fatal clinical outcomes, including UTI or sepsis. Therefore, urine and blood represent relevant clinical matrices for *Kp* diagnostics. Using our reporter phage rTUN1::*nLuc* on both *Kp*-spiked (1×10^6^ CFUs/well) urine and blood samples, respectively, distinct luminescence peaks could be measured. Conversely, no matrix associated increase in background luminescence was detected (Figure 4C). While the results for spiked urine samples were almost identical to those measured in spiked growth medium, the luminescence peaks retrieved from spiked blood samples appeared to be time-shifted by 2 h now correlating with the results of an initial *Kp* concentration of 1×10^4^ CFUs/well. This can most easily be explained by a loss of 10^2^ bacteria per ml during serum collection from spiked blood.

### rTUN1::*nLuc* is highly specific for *Kp* K64 strains

Recently, we described that TUN1 WT exclusively infects *Kp* K64 strains (Eckstein et al., 2021). In order to verify the host specificity of rTUN1::*nLuc*, we analyzed the luminescence development in liquid culture testing four additional *Kp* K64 strains and five *Kp* strains with other K-types (K55, K52, K27, K25 and K17) in the presence of the reporter phage. Positive signals (> 10^3^ RLU) were obtained only from samples containing *Kp* K64 strains indicating that rTUN1::*nLuc* features the same specificity for *Kp* K64 strains as the WT phage (Figure 4D).

### rTUN1::*nLuc* enables rapid, real-time antibiotic susceptibility testing of *Kp*

The main challenge for treatment of *Kp* infections is the increasing number of MDR *Kp* strains. For instance, *Kp* 7984, the strain used in this work, has already been shown to be resistant to the first-line AB ertapenem (Kollenda et al., 2019). Testing of antibiotic susceptibilities of *Kp* strains derived from patient samples is therefore crucial to ensure successful therapy. However, these tests require a pure culture of the infection-causing strain and often contain long incubation steps. To accelerate this process of selecting the adequate ABs for an efficient therapy, we tested whether rTUN1::*nLuc* is also suitable for the real-time, rapid AB susceptibility testing. For this, we used the experimental settings of the luminescent reporter phage assay (10^6^ CFUs/well *Kp* 7984 + 10^2^ PFUs/well rTUN1::*nLuc*, shown in Fig. 4A) and supplemented the growth medium with various ABs of different classes using their minimal inhibitory concentration (MIC) at breakpoints determined for *Enterobacteriaceae* (Supplementary Table S1). We observed no increase in reporter phage associated luminescence in the presence of gentamicin (Gent), chloramphenicol (Cmp), imipenem (Imp), meropenem (Mer) and amoxicillin/clavulanic acid (Amc) (Figure 5A). In contrast, we detected a strong luminescence increase resulting in a distinct peak after 2 hours, comparable to that of the *Kp* 7984 + rTUN1::*nLuc*-control without ABs, when levofloxacin (Lev), ceftazidim (Caz), streptomycin (Str), tigecyclin (Tgc+), ciprofloxacin (Cip) and trimethoprim/sulfamethoxazol (T/S) had been added (Figure 5B). Growth experiments of *Kp* 7984 (without phage) in the presence of ABs further revealed that Gent, Cmp, Imp, Mer and Amc inhibited bacterial growth whereas *Kp* 7984 growth was not affected by the presence of Caz, Str, Tgc+, Cip and T/S (Supplementary Figure S5). Thus, *Kp* 7984 can be considered sensitive to Gent, Cmp, Imp, Mer and Amc and resistant to Caz, Str, Tgc+, Cip and T/S. In contrast to time consuming classical growth experiments and MIC testing, our luminescent reporter phage assay using rTUN1::*nLuc* not only enables rapid and sensitive detection of *Kp* K64 strains directly from clinical samples, but also allows for simultaneous, real-time antibiotic susceptibility testing within just a few hours.

**Figure 5.**
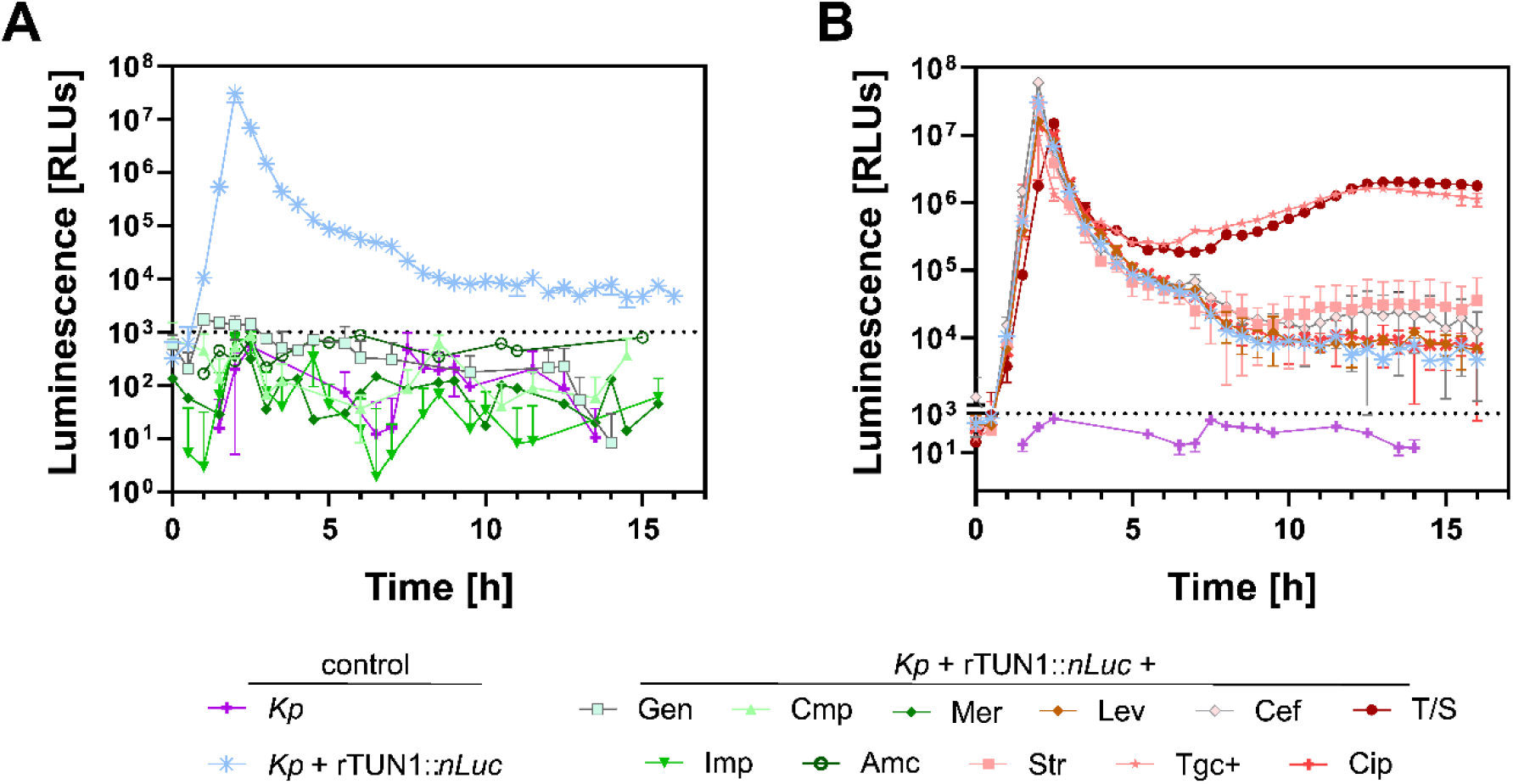
Antibiotic susceptibility testing of *Kp* using rTUN1:: *nLuc*. Luminescence measurements of wells containing rTUN1::*nLuc*-infected *Kp* 7984 (1×10^6^ CFUs + 1×10^2^ PFUs per well) in the presence of different antibiotics. *Kp* without phage (purple) and *Kp* + rTUN1::*nLuc* in pure LB (blue) served as controls **(A)** Despite the positive control no luminescence peak was detected for wells containing *Kp* + rTUN1::*nLuc* in gentamicin (Gen), chloramphenicol (Cmp), imipenem (Imp), meropenem (Mer) or amoxicillin/ clavulanic (Amc). **(B)** Distinct luminescence peaks were generated in samples containing *Kp* + rTUN1::*nLuc* with levofloxacin (Lev), ceftazidime (Caz), streptomycin (Str), tigecyclin (Tgc+), ciprofloxacin (Cip) or trimethoprim/ sulfamethoxazole (T/S). Error bars represent three independent biological replicates.

## Discussion

The enormous increase of antibiotic resistances in bacteria has been estimated to lead to an excess mortality of 10 million persons per year by 2050, mostly caused by nosocomial infections (Alebachew Woldu, 2016; Borer et al., 2009). The multiple antibiotic resistance crisis constitutes a tremendous threat to global health and has been declared a global health emergency (Toner et al., 2015) – a view supported by WHO. Rapid pathogen detection and antibiotic resistance screening allows non-empirical therapy and saves lives. Established methods for *Kp* detection are time consuming and necessitate empirical antibiotic therapy in urgent cases (Davenport et al., 2017) with more unfavorable outcomes. Here, we demonstrated that reporter phage-based diagnostics could be a promising alternative, as integration of reporter genes into highly specific phages enable real-time detection of phage replication and thus, living host cells. Besides fluorescent proteins (Jain et al., 2012) and hydrolyzing enzymes such as β-galactosidase (Chen et al., 2017; Chen et al., 2017)), luciferases can be used as reporters (Hinkley et al., 2018; Loessner et al., 1996b; Meile, Sarbach, et al., 2020; Nguyen et al., 2017).

Since the discovery of luciferin-luciferase systems almost a century ago (McElroy & Ballentine, 1944), several luciferase enzymes have become available to researchers. The latest commercially available luciferase is the NanoLuc^®^ enzyme (nLuc) which has several advantages compared to other luciferase systems: It features a relatively small size of only 19.1 kDa (compared to e.g. the Renilla or Firefly luciferase with 36 kDa or 61 kDa, respectively), fast substrate turnover rates – which leads to generation of a high-intensity luminescent signal - and exhibits high stability under various buffer conditions (England et al., 2016; Hall et al., 2012). Therefore, nLuc was selected as reporter candidate for this study.

Inserting reporter genes, or any additional nucleotide sequence, into phage genomes can be challenging. As DNA length and capsid size and thus internal pressure in the capsid correlate, (Nurmemmedov et al., 2007) the DNA encapsulation capacity of most phages is limited. Hence, integration of additional genes can negatively affect capsid stability and therefore lead to severe impairments in phage assembly or replication (Grayson et al., 2006; Nurmemmedov et al., 2012). DNA packaging of tailed phages relies on the recognition of specific sequences (*cos-* or *pac-*sites) by the terminase protein followed by DNA cleavage. Depending on the phage type, this cleavage can be either sequence-specific (e.g. T7 or λ) or nonspecific (e.g. P22 or SPP1). For the latter, the cleavage does not occur at the end of the phage genome (so-called termination cleavage) but follows the mechanism of headful packaging where nonspecific DNA cleavage by TerL (terminase large subunit) is induced by increased capsid pressure. As a result, phages using the headful system often tolerate a genome size of up to 110% while the others will not (Casjens & Gilcrease, 2009). Since TerL initiates DNA packaging and cleavage, phylogenetic analysis of the *terl* gene can be used to predict a phage’s DNA packaging system. For TUN1, results of this analysis suggested that the phage belongs to the “T7-like group with directed terminal repeats” and not to the “P22-like headful group”. Therefore, the fact that our *nLuc* reporter construct could not be inserted into the full-length TUN1 WT genome can most likely be explained by its sequence-specific DNA packaging mechanism in combination with a genome size that has already reached the maximum capsid capacity.

It has already been described, that insertion of additional genetic information into the genomes of other *Podoviridiae* or *Autographiviridae* requires preceded genome reduction in order to overcome the problem of limited capsid capacity (Kim et al., 2014; Pires et al., 2021). In this work, we were able to minimize the TUN1 genome by up to 4.3 % (TUN1 Δhpgc123) and, at the same time, demonstrated that the hypothetical genes *gp2-4*, *gp7-8* and *gp10-11* are not essential for a successful lytic cycle of TUN1. Using the genome-reduced TUN1 Δhpgc2 as a template, insertion of the RBS-*nLuc* reporter construct was successful. The resulting, luminescent reporter phage, rTUN1::*nLuc*, enabled specific and sensitive detection of *Kp* K64 cells exhibiting a limit of detection (LOD) of only CFU per well.

To our knowledge, there are only two other *Kp*-specific nLuc-based reporter phages, Mcoc and 8M7, that specifically detect *Kp* K21 cells with a comparably low LOD of only 10 CFUs/well and 100 bacteria per 100 mg feces, respectively (Zelcbuch et al., 2021). Other *nLuc-*based reporter phage assays exhibited similar sensitivity. For instance, Pulkkinen et al constructed a T7-*nLuc* phage, which was able to detect 47 *E. coli* cells per well (Pulkkinen et al., 2019). Further, ΦV10nluc phage-based *E. coli* detection exhibited an even lower LOD of only 5 CFUs/well. These assays, however, were not tested on clinically relevant matrices. In contrast, our novel rTUN1::*nLuc* based reporter phage assay enables rapid detection of *Kp* cells from blood and urine samples exhibiting no matrix effects such as luciferase quenching or unspecific increase in background luminescence. While the assay detected *Kp* K64 in urine with the same sensitivity as pure bacterial culture, luminescence peaks retrieved from spiked blood samples indicated a loss of bacterial cells during sample preparation. This was most probably caused by the washing steps during serum collection rather than by blood associated matrix effects as comparable studies have shown that similar optimization steps (serum separation and washing) of spiked blood samples resulted in a 20-50% reduction in bacterial concentration (Moses et al., 2021; Sergueev et al., 2017).

The current cutoff for diagnosis of UTIs, one of the most common diseases caused by *Kp*, is 1×10^5^ CFUs/ml urine (Becknell et al., 2015). Using our novel reporter phage assay, this minimal pathological *Kp* load can be detected within four hours directly from the patient’s urine. Compared to goldstandard methods like PCR, rTUN1::*nLuc* only detects viable *Kp* cells, which can be crucial, e.g., for monitoring therapy success where differentiation between living and dead cells is essential.

In addition to that, our approach allows for rapid, real-time assessment of antibiotic resistances (real-time antibiogram) and thus, facilitates treatment of *Kp* K64 infections. Here, susceptibility against therapeutically relevant antibiotics can be tested simultaneously to initial *Kp* detection from clinical samples providing vital information for assessing treatment options in a few hours (depending on the *Kp* load in the sample). In case of an infection with a MDR *Kp* K64 strain with no promising treatment options left, rTUN1::*nLuc* can also function as a companion phage for phage therapy, as positive luminescence readout from reporter phage assays also indicate infectivity of the wildtype TUN1 and thus, qualifies for phage therapy.

In conclusion, our results clearly demonstrate the enormous diagnostic capabilities of reporter phages. However, for broad applicability directly from clinical samples, such as urine from patients suffering from UTIs, reporter phages need to be constructed not only for all locally occurring *Kp* K types, but for all clinically relevant pathogens. This would then allow not only the rapid identification of the infection-causing bacterial strain from unknown samples within a few hours, but also the simultaneous generation of a real-time antibiogram of the infection-causing strain as well as the selection of potential candidates for phage therapy.

## Acknowledgements

We are grateful to Prof. Dr. Ben Moussa from the Military Hospital of Instruction in Tunis, for providing the waste water samples from which we could initially isolate the bacteriophage TUN1, and Daniela Friese for technical assistance.

## Funding

This study was funded by the Medical Biological Defense Research Program of the Bundeswehr Medical Service and supported by supported by the Enable and Enhance Initiative of the federal government of Germany (Ertüchtigungsinitiative der Deutschen Bundesregierung, grant no. OR12-370.43 ERT TUN IMB).

## Authors’ contribution

Conceptualization: S.E., J.J.B and P.B.; investigation: R.R., P.B. and S.E.; methodology: S.E. and P.B.; formal analysis and validation: P.B., R.R. and S.E.; visualization: S.E.; writing—original draft preparation: S.E. and P.B.; writing—review and editing: P.B., J.J.B. and S.E.; supervision and project administration: JJ.B. and S.E. All authors have read and agreed to the published version of the manuscript.

## Notes

### Competing Interest Statement

The authors have declared no competing interest.

### Summary of Updates

text

