## Supplementary material for "Recombinant reporter phage rTUN1::*nLuc* enables rapid detection and real-time antibiotic susceptibility testing of *Klebsiella pneumoniae* K64 strains": Suppl. Material (from another version of this preprint: version 1, 2022-08-19)

### Supplementary Figures

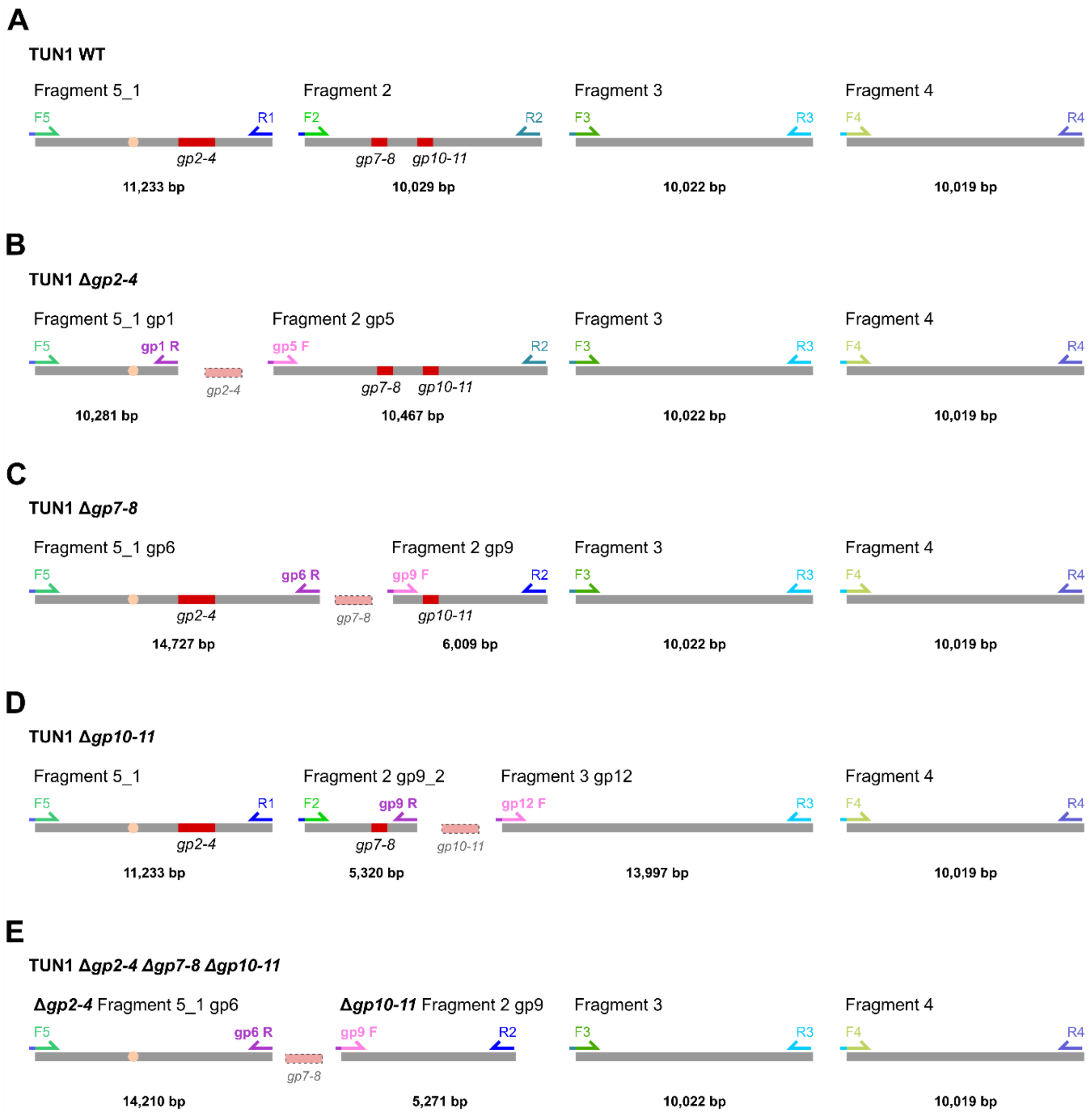


**Supplementary Figure S1. Schematic overview of synthetic TUN1 fragments used for *in vitro* DNA assembly.** Amplicons used to assemble TUN1 WT (**A**), TUN1 Δ*gp2-4* (**B**), TUN1 Δ*gp7-8* (**C**), TUN1 Δ*gp10-11* (**D**) and TUN1 Δ*gp2-4* Δ*gp7-8* Δ*gp10-11* (**E**) *in vitro*. Gene clusters of interest (for deletion) are marked in red. Deleted gene clusters are marked with a pitched line. One-sided arrows depict primers used for fragment amplification. Orange dots represents genome circularization. If not stated differently, gDNA of TUN1 WT was used as PCR template.

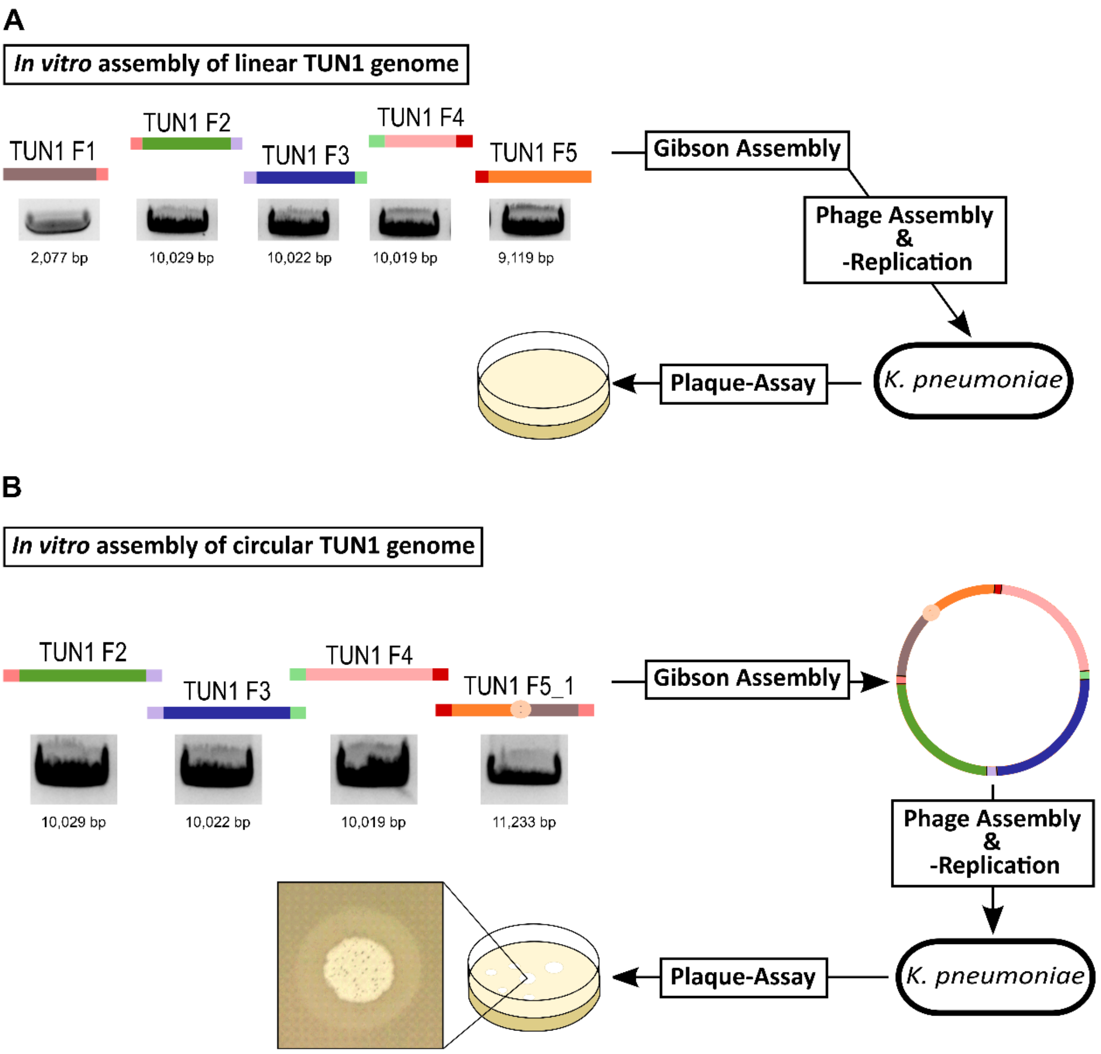


**Supplementary Figure 2. Schematic overview of different approached for *in vitro* assembly and rebooting of synthetic TUN1 WT genome.** For assembly and rebooting of TUN1 WT all synthetic genome fragments were amplified with overlapping ends between consecutive segments (depicted in matching colors). Subsequently, plaque assays on *Kp* were performed using transformation supernatants. **A)** *In vitro* DNA assembly of linear TUN1 genome. **B)** *In vitro* DNA assembly of circularized TUN1 genome. Plaques, and therefore functional phages, were only obtained when circularized synthetic genomes were used for phage rebooting.

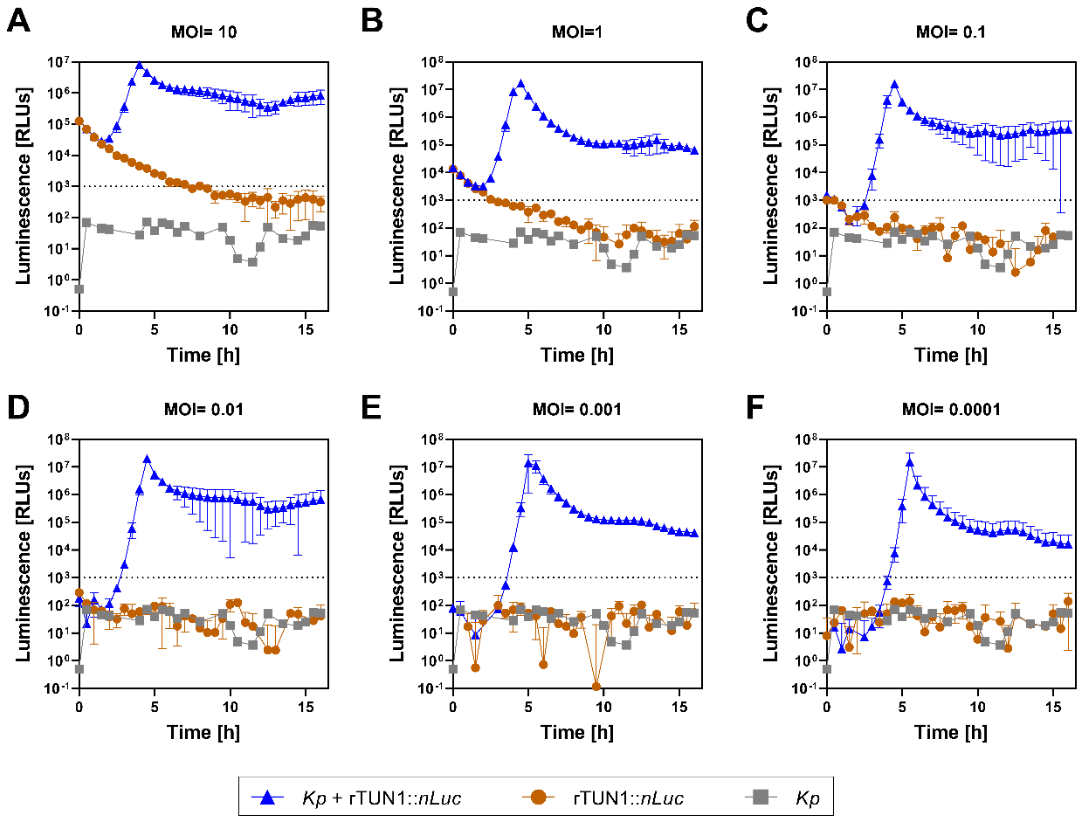


**Supplementary Figure 3. Effects of reporterphage titration on luminescence signal.** Luminescence signal derived from $1x10^4$ CFUs/well *Kp* 7984 mixed with different rTUN1::*nLuc* concentrations per well (blue triangles): **A)** $1x10^5$ PFUs (MOI= 10) **B)** $1x10^4$ PFUs (MOI= 1) **C)** $1x10^3$ PFUs (MOI= 0.1) **D)** $1x10^2$ PFUs (MOI= 0.01) **E)** $1x10^1$ PFUs (MOI= 0.001) **F)** $1x10^0$ PFUs (MOI= 0.0001)

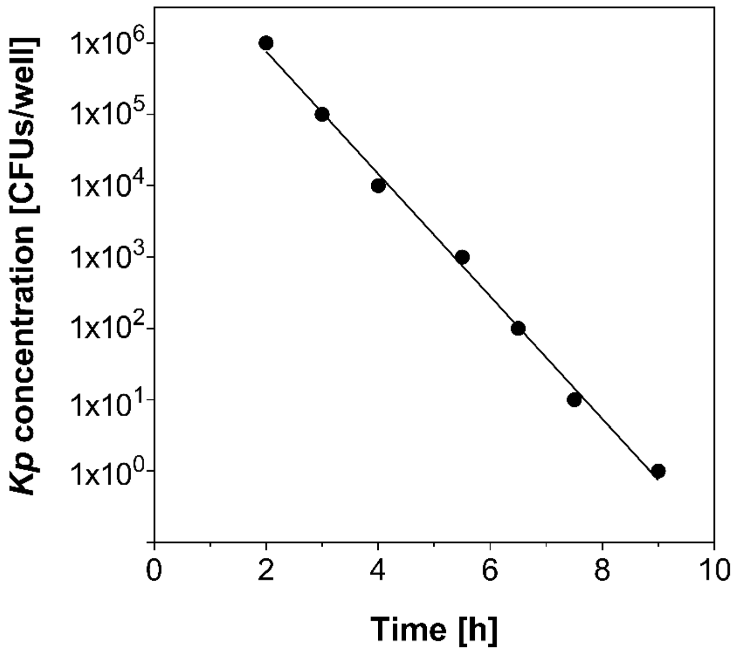


**Supplementary Figure 4. Standard curve for *Kp* concentration determination.** Timing of luminescence peak as a function of *Kp* concentration to allow *Kp*-load determination in unknown samples.

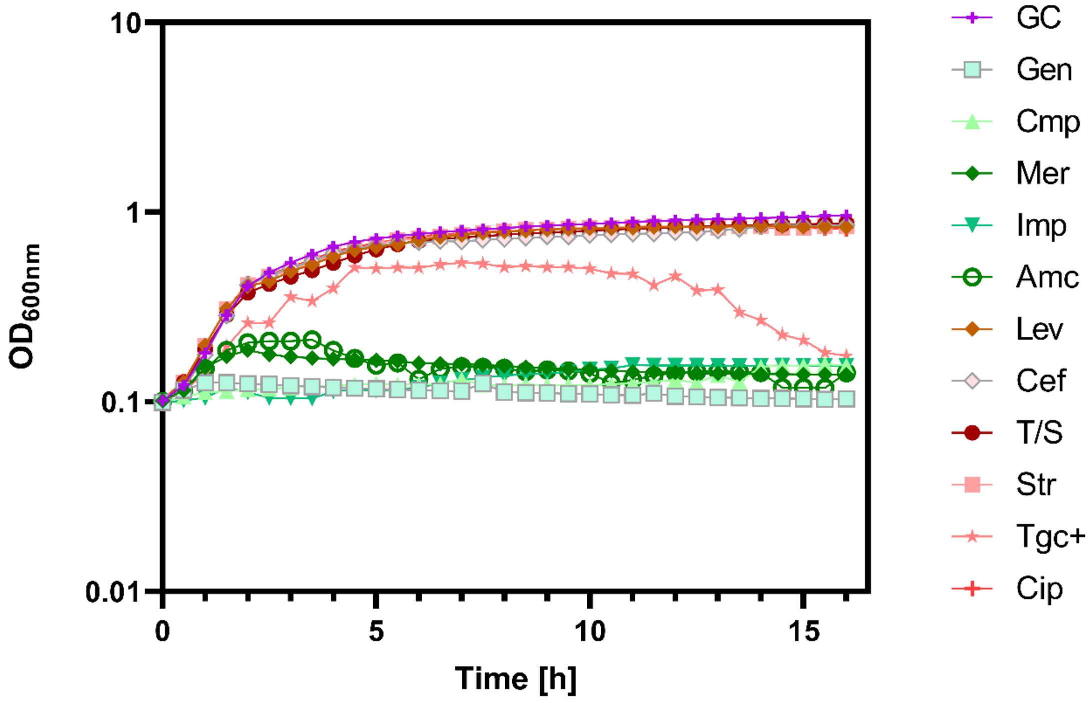


**Supplementary Figure S5. Growth curve of Kp 7984 with various antibiotics.**

### Supplementary Tables

**Supplementary Table S1. Antibiotics used for susceptibility screening of *Kp* 7984 together with rTUN1::*nLuc.*** Minimal inhibition concentrations (MICs) breakpoints referring to antibiotic resistance were adapted from "The European Committee on Antimicrobial Susceptibility Testing. Breakpoint tables for interpretation of MICs and zone diameters. Version 12.0, 2022. http://www.eucast.org"

| Antibiotics | MIC breakpoints R [mg/L] |
|---|---|
| **Penicillins** | |
| Amoxicillin/clavulanic acid (uncomplicated UTI only) | > 32 /16 |
| **Chepalosporins** | |
| Ceftazidime | > 4 |
| **Carbapenems** | |
| Imipenem | > 4 |
| Meropenem | > 2 |
| **Fluoroquinolones** | |
| Moxifloxacin | > 0.25 |
| Ciprofloxacin | > 0.5 |
| Levofloxacin | > 1 |
| **Fluoroquinolones** | |
| Gentamicin | > 2 |
| Streptomycin | > 1 |
| **Tetracyclines** | |
| Tigecycline | > 0.5 |
| **Other** | |
| Trimethoprim/sulfamethoxazol | > 4/76 |
